# Comparing Random and Natural RNA Boltzmann Ensembles

**DOI:** 10.64898/2026.03.31.715513

**Authors:** Hiba Khan, Paula Garcia-Galindo, Sebastian Ahnert, Kamal Dingle

## Abstract

A *morphospace* is an abstract space of theoretically possible biological traits, shapes, or property values. It is interesting to explore which parts of a morphospace life occupies, as compared to those parts which could be occupied, but are not. Comparing random and natural non-coding (nc) RNA secondary structures is an established approach to studying morphospace occupation for RNA structures. Most earlier studies have focused on the minimum free energy (MFE) structure, while relatively few have looked at the Boltzmann distribution, describing the ensemble of energetically suboptimal RNA folds. These suboptimal structures may have important roles and functions, and hence should be examined carefully. Here we compare random and natural ncRNA in terms of their Boltzmann distributions, finding that natural RNA tend to have very similar profiles to random RNA, with the main difference being that natural RNA are slightly more energetically stable, except for very short sequences (20 to 30 nucleotides) which appear to be equally or slightly *less* stable. We infer that natural ncRNA and random RNA occupy similar parts of the morphospace, indicating that the biophysics of the genotype-phenotype map largely determines the ensemble properties of ncRNA.

## I. INTRODUCTION

A fundamental aim of biophysics is to understand the varied structures present in organisms. In addition to invoking natural selection to explain biological traits and properties [1, 2], many patterns and structures can be rationalized through physical, mathematical, and mechanical arguments [3, 4], as well as by development [5], mutation biases [6], self-organization [7, 8], or other factors [9].

Following this mathematical perspective, it is useful to think of a *morphospace* [10–13], i.e., an abstract mathematical space of biological traits, values, or properties. Given this space, we can then ask questions like, which parts of the morphospace are occupied during the course of biological evolution [14], and why? Although easy to ask, answering these questions is often challenging due to the difficulty of untangling the various factors and their contributions over time.

One system where progress can been made is the mapping of RNA sequences to RNA secondary structures, which has been extensively studied as a toy model genotype-phenotype map due to being computationally tractable yet biologically relevant and realistic [15–22]. Because RNA has 4 possible nucleotides per sequence base, there are 4^*L*^ different possible genotype sequences of length *L*. Using computational methods, secondary structures (i.e., nucleotide bonding patterns) can be predicted from sequences [23], which gives the phenotypes. The number of different secondary structures grows as *∼* 1.76^*L*^ [19]. By enumerating or sampling genotypes, the theoretical morphospace of possible RNA secondary structures can be determined, along with many quantities such as the total number of possible phenotypes, and the distribution of genotypes per phenotype [19, 24]. In short, the theoretical morphospace can be well characterized for this map.

Characterizing morphospace occupation of natural RNA is also relatively tractable due to the availability of natural sequence data found in bioinformatics databases [25]. Statistical properties of the morphospace of these natural sequences can be compared to properties of randomly generated sequences, where the latter can act as a null-model, and as a basis from which to infer the action of selection (or lack therefore). Interestingly, natural non-coding RNA shows similar statistical properties to those of randomly generated RNA sequences [8, 17, 26–28], e.g. in terms of the numbers of sequences per structure. Dingle et al. [19] advanced the analysis of RNA morphospaces by inferring not only the properties of RNA generated from random sequences, but also those of RNA from the full morphospace of possible secondary structures. The full morphospace being the set of all possible RNA structures (with at least one genotype). This advancement showed that not only are random and natural structures similar to each other, but that they are both very different from typical structures from the full space. Later, it was shown [20, 29] that not only secondary structure, but the overall coarse-grained shapes and motifs are similar too, and that natural RNA tend to be only those which are most easily found in the sequence space, i.e., those structures which can accommodate the highest number of possible sequences per structure. Moreover, the frequencies of shapes found in nature correlate strongly with the frequencies observed in samples of purely random sequences. Recently, von der Dunk et al. [30] made a detailed study of different natural RNA and lengths, confirming these similarities in morphospace occupation for larger RNA of lengths *L* = 300.

These observations of the similarity between natural and random non-coding RNA are by no means obvious or trivial, because a foundational tenet of molecular biology is the close link between structure and function [31–33] (e.g., tRNA, rRNA, ribozymes, riboswitches). This tenet suggests that RNA would have tuned its specific structure through natural selection to meet the needs of a biological function, which would not necessarily correlate with the properties of random sequences. Indeed, it is known that RNA secondary structure is under selection [34–36], in addition to the sequence motifs themselves which must also be under selection for e.g. binding sites. Additionally, as pointed out by von der Dunk et al. [30], RNA molecules are highly evolvable, and so are able to adopt a variety of different shapes when under selection. That is to say, it is not at all impossible for evolution to reach non-random structures. Hence the close similarity between natural and random RNA remains somewhat perplexing.

In biology it is common to think of the minimum free energy (MFE) structure as ‘the’ secondary structure of an RNA molecule, but in reality this is a simplification; a given sequence is dynamically switching between different Boltzmann distributed structures, because of thermal fluctuations at the molecular scale [22, 32, 37–41]. By definition, a sequence will occupy the MFE configuration more frequently than any other structure, but depending on the energy levels of the other competing sub-optimal structures, the time spent in other than the MFE structure may be substantial. Additionally, suboptimal structures in biology can impact function [41–47], even to the extent of a proposed “RNA-ensemble–function paradigm” [48], and there is evidence that suboptimal structures are under selection [49].

The earlier comparisons of natural and random RNA (see above) have almost entirely focused on MFE structures. Notable exceptions include the work of Miklos et al. [50] who studied Boltzmann distributions for RNA, including comparing natural and random sequences. They compared the free energy values of natural miRNA, 5S rRNA, and tRNA (from *E*.*coli*). The main findings were that the distribution of energies for miRNA (which are very short with ≈ 22 nucleotides) was substantially different from random sequences (implying the Boltzmann probabilities differ, too), while for 5S rRNA and tRNA the distributions of energy values were similar. Chan and Dong [51] also made a brief related study to Miklos et al., reporting that the Boltzmann distribution for a very small sample of mRNA was similar to random sequences. On the other hand, they reported the statistical distinguishability of small samples of natural and random non-coding RNA, especially miRNAs.

Here we extend the investigation of the morphospace occupation in natural non-coding RNA by analysing several metrics of suboptimal structures, i.e. the energetically less favorable structures, for a diverse RNA data set of different lengths.

## II. HOW SIMILAR ARE RANDOM AND NATURAL SUBOPTIMAL STRUCTURE ENSEMBLES?

### A. Natural RNA data and prediction algorithms

We will compare random and natural non-coding RNA. For the natural RNA, we downloaded all available sequences of lengths *L* = 20, 30, 45, 60, 100, 150 from the RNAcentral database [25] (Methods, Appendix A). The number of natural sequences used in our analysis is 12,438 (*L* = 20), 47,975 (*L* = 30), 13,541 (*L* = 45), 20,897 (*L* = 60), 25,533 (*L* = 100), 15,494 (*L* = 150).

For the random sequence data, a pseudo-random number generator was employed to make random sequences of the same lengths as the natural data. Each nucleotide A, T(U), C, G was assigned a 25% chance of appearance.

We study *L* = 20, 30, 45, 60, 100, 150 because these lengths are computationally tractable, similar to the lengths which have been used in earlier studies [19, 28], structure prediction algorithms are more accurate (and hence reliable) for shorter sequences [52, 53], and because structure can be important even for very small RNA of *L* ≈ 45 [54] and *L* ≈ 20 [55].

The MFE secondary structures and suboptimal structures were predicted computationally via energy minimization using the well-known RNA Vienna package [23], with default parameters (Methods, Appendix A). The algorithm gives predicted energy values for the MFE and suboptimal structures to within a threshold given in kcal/mol. Throughout this work, we use 10.0 kcal/mol as the threshold for *L* ≤ 100, and 5.0 kcal/mol for *L* = 150. Using a larger threshold may yield more accurate results by incorporating a greater variety of suboptimal structures, but there are exponentially many different such structures which increases computational resource requirements. See Appendix C for more discussion around the choice of energy threshold cutoff.

RNA sequences populate the ensemble (consisting of the MFE and suboptimal structures) according to the Boltzmann distribution, given by

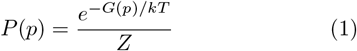

where *G*(*p*) is the free energy of a given structure *p* within the ensemble, *Z* is the partition function, *k* is the Boltzmann constant, and *T* is the absolute temperature in Kelvin. The fraction of time spent in a given secondary structure *p* with free energy *G*(*p*) is given by *P* (*p*) [56]. It follows that if the MFE structure has a relatively low energy value *G*(*p*), while the other structures in the ensemble have higher energies, then the MFE will dominate the ensemble. On the other hand, if the energy of the MFE structure is not relatively very low, and other structures in the ensemble have similar energies, then the other structures in the ensemble may dominate.

Using the natural and random RNA sequences, we quantitatively compare the secondary structures and Boltzmann probability distributions in several ways. We are not primarily concerned with small but statistically significant differences; rather, in line with the perspective of morphospace occupation, we ask for some biophysically plausible range of values, how similar are natural and random distributions and/or other statistics?

Figure 1 visually depicts the key ideas of our work, including morphospaces, sequences, shapes, and Boltzmann ensembles of suboptimal structures.

**FIG. 1.**
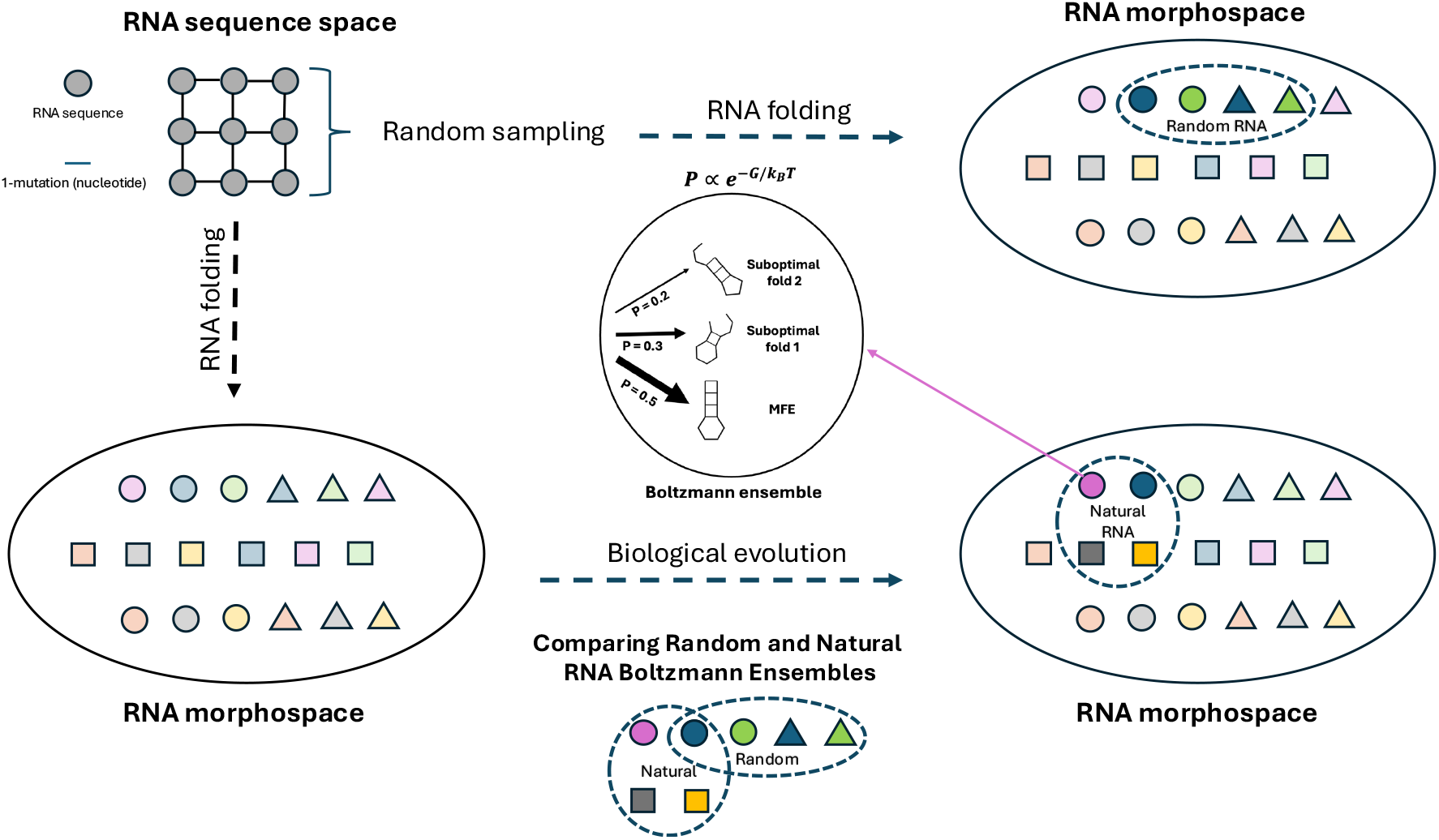
Schematic of random and natural RNA Boltzmann ensembles in the RNA morphospace, and their comparison. In this work, the RNA morphospace describes the set of all possible Boltzmann ensembles for an RNA of sequence length *L*. It is constructed by folding RNA sequences in sequence space, using ViennaRNA software [23], and taking from these the set of different possible Boltzmann ensembles. The shapes and colours are used to illustrate the different structures in morphospace, in this case, the different Boltzmann ensembles. An example of a Boltzmann ensemble for RNA of length *L* = 12 is shown in the **(middle)**, where the MFE is the minimum free energy structure, and the two others are the suboptimal structures, ranked by their probability, which is proportional to the Boltzmann distribution. **(top)** An extremely simple illustration of RNA sequence space, which is composed of all RNA sequences of a certain length *L*. Each sequence is connected to each other through 1-point mutations (change of a nucleotide base letter). The sequence space is randomly sampled and then the sampled sequences are folded to create the set of Boltzmann ensembles which determine the random RNA occupation in the RNA morphospace. **(bottom)** From the total RNA morphospace, the course of biological evolution has defined a set occupation, which we call the natural RNA Boltzmann ensembles. We can infer the natural RNA morphospace occupation by folding the non-coding RNA sequences found in biological experimental databases. The comparison between the random and natural RNA Boltzmann ensembles can show to what extent the selection process in biological evolution or the statistical properties of sequence space, have shaped natural RNA structures.

### B. Rank plots of Boltzmann probabilities

We begin the comparison of natural and random ensembles by visualizing the Boltzmann probabilities. Figure 2 shows rank plots of the mean Boltzmann probabilities for the MFE and first 7 suboptimal structures. Rank 1 corresponds to the MFE structure, rank 2 to the first suboptimal structure, etc. As can be seen for all lengths, typically the MFE structure is occupied for much less than 100% of the time, and the suboptimal structures account for more than 50% of the probability, indicating the importance of studying suboptimal structures in addition to the MFE. The standard deviation is large in all cases, implying considerable variation in this quantity depending on the sequence.

**FIG. 2.**
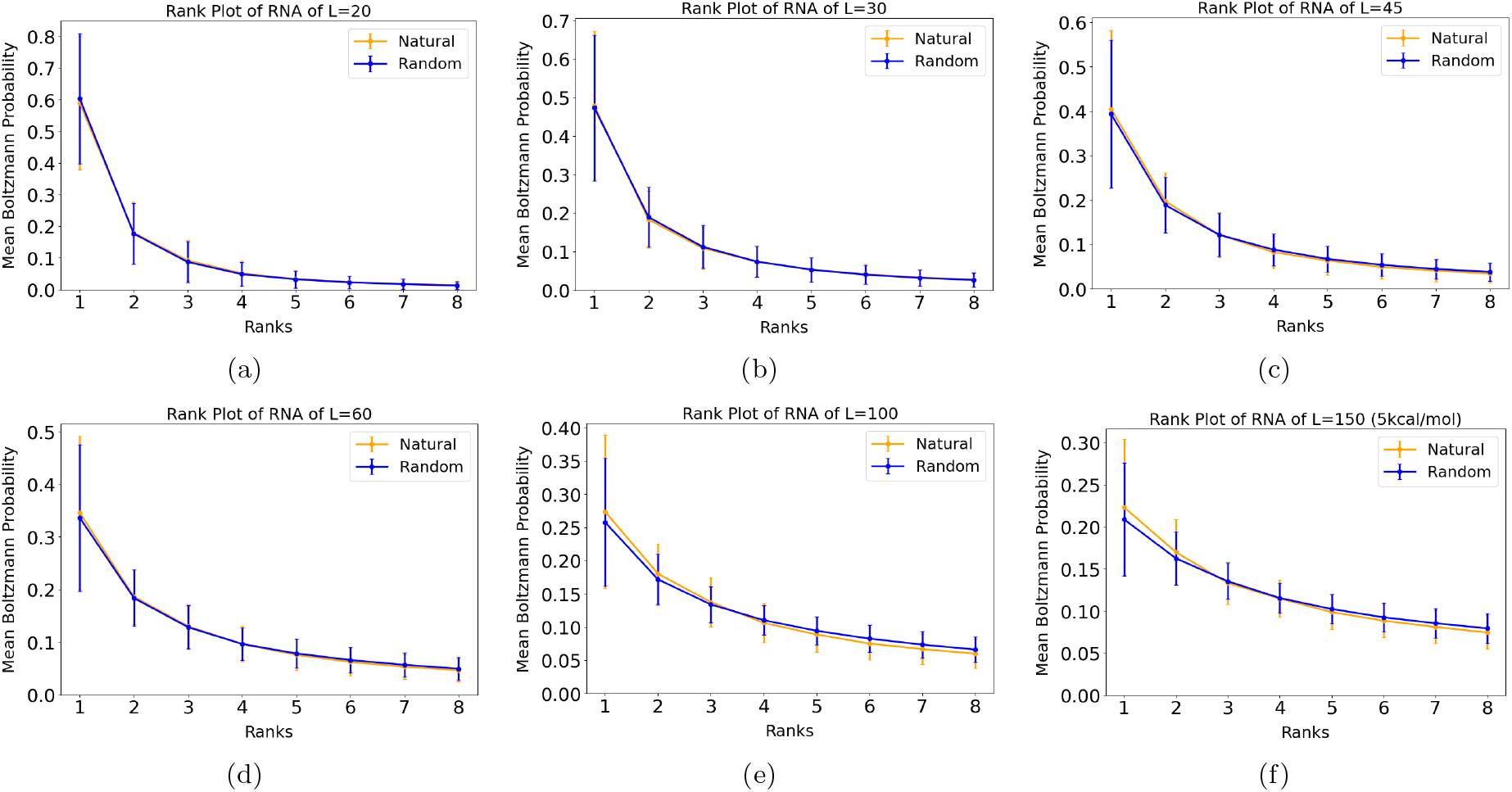
Mean Boltzmann probability rank plots. Rank 1 is the minimum free energy (MFE) structure within the ensemble, rank 2 is the first suboptimal structure, rank 3 the second suboptimal structure, etc. (a) *L* = 20, (b) *L* = 30, (c) *L* = 45, (d) *L* = 60, (e) *L* = 100, and (f) *L* = 150. In all cases, the probability of rank 1 is much less than 1.0, indicating the importance of studying the suboptimal structures which account for a substantial amount of the probability mass. Natural and random distributions are very similar for all lengths *L*. For each length *L*, data from 1000 random and 1000 natural sequences were used.

For some lengths (*L* = 100, 150), the mean Boltzmann probability of the MFE is slightly higher for natural than for random RNA (as reported earlier by [50] for three types of natural RNA). However, we note that, overall, the natural and random curves are similar. To quantify the similarity, we can compare natural and random mean Boltzmann probability values of each of the 8 ranked structures in Figure 2, for each length *L*. With *L* = 20, the natural mean Boltzmann probability values as a percentage of the random mean Boltzmann probability values are 98%, 101%, 106%, 104%, 103%, 100%, 100%, 100%. These 8 mean values have a mean of 101%. Because 101% is very close to 100%, this shows that the mean Boltzmann probabilities of the first 8 ranked structures of the natural and random structures are generally very similar. For *L* = 30, the mean value is also 101%; for *L* = 45, the mean value is 96%; for *L* = 60, the mean value is 98%; for *L* = 100, the mean value is 97%; and for *L* = 150, the mean value (using 5.0 kcal/mol) is 99%. Hence, for each length, the mean is close to 100% and so the natural and random mean Boltzmann probabilities for the first 8 ranked structures are close.

### C. Combined shapes

Traditional dot-bracket representations of RNA structures depict all bonds. It can be beneficial to study more abstract or coarse grained RNA shapes, which do not depict every detail, but instead capture the overall shape [57]. This is because presumably the overall shapes or motifs of non-coding RNA are more relevant to its function, than every detail of the structure. One practical advantage of using abstract shapes is that it is easier to compute reliable frequency estimates for structures, due to the increased ratio of sequences per shape (as compared to sequences per dot-bracket structure) [20]. The abstract shapes can be computed for 5 different levels, with Level 1 retaining the most detail, and Level 5 being the most abstract [57]. For example, the dot-bracket secondary structure (((((…..)))))…((…)) can be abstracted to just two stems, written in abstract notation as [][].

Taking the abstract shape for the MFE structure, and abstract shapes of the other suboptimal structures, we can form a “combined phenotype” by concatenating these. For example, if a sequence has abstract shape [[]] as the MFE, [] as the first suboptimal shape, and [][] as the second suboptimal shape, then the combined phenotype would be [[]]*[]*[][], where the asterisk “*” indicates concatenation. Using this procedure, we can compare the frequency of combined phenotypes in random and natural data (using abstract Level 5 as in [20]). The benefit is that this gives a way to compare not just the Boltzmann probabilities but the shapes themselves.

In Figure 3, we show the frequencies of combined shapes for natural and random RNA, where the MFE shape and the first, second, third, and fourth suboptimal shapes are combined (so that 5 shapes are combined together). The linear correlations of log-frequencies are high at 0.98, 0.98, 0.92, 0.89, 0.84, 0.75, and all *p*-values are statistically significant (*p*-values ≪ 0.05) implying that the natural and random are very similar in terms of suboptimal shapes. Note that only combined shapes with non-zero frequency in both the natural and random data were plotted (log frequency is undefined when frequency is 0).

**FIG. 3.**
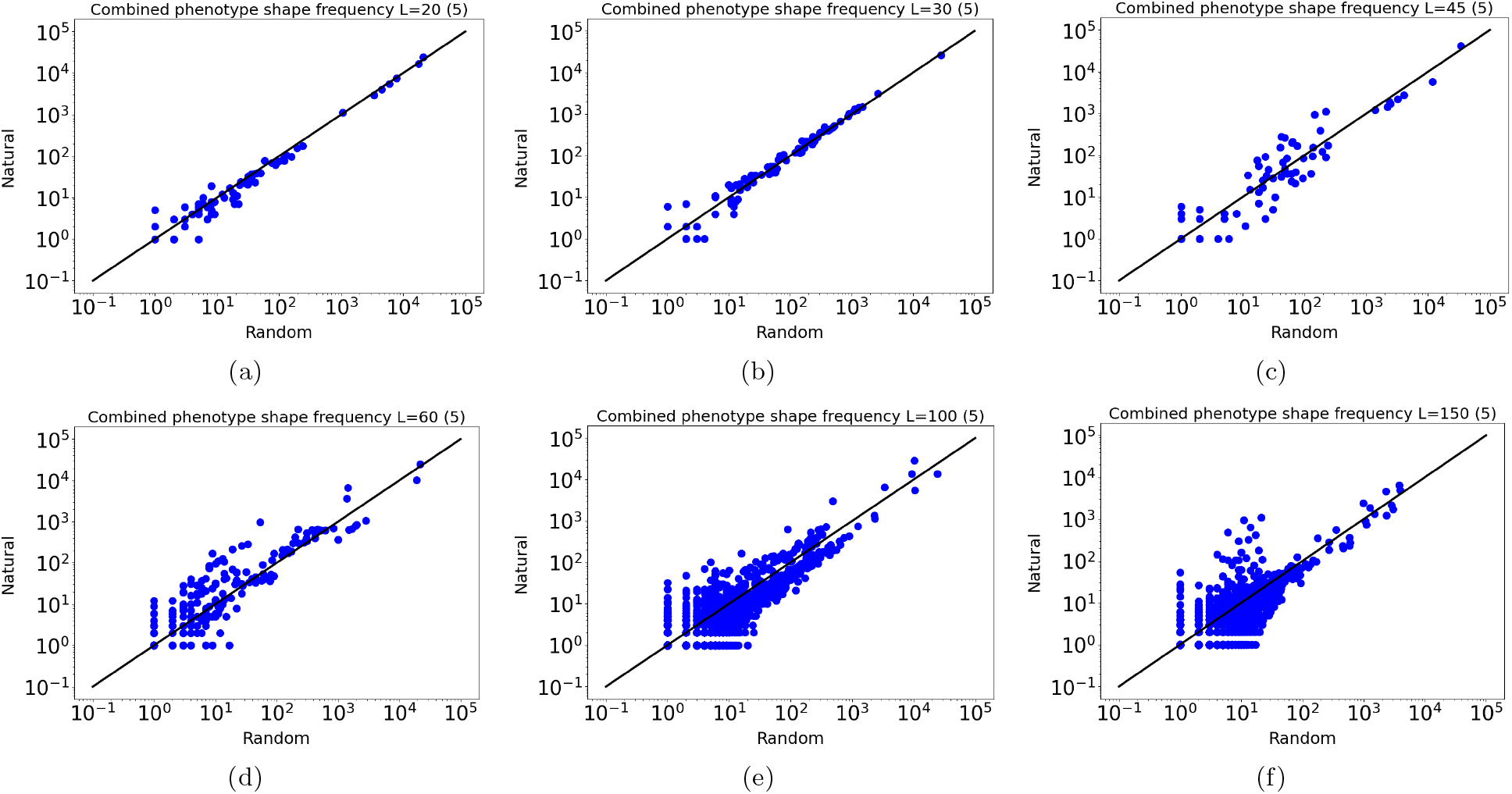
Linear correlations in logarithm of combined shape frequencies. (a) *L* = 20, (b) *L* = 30, (c) *L* = 45, (d) *L* = 60, (e) *L* = 100, and (f) *L* = 150. Five shapes are concatenated together (the MFE shape and the first four suboptimal shapes). The correlations are high for all lengths.

As *L* increases the number of different shapes increases, so that the correlations become more noisy due to the lower frequency count numbers of sequences per shapes. This helps to understand the decreasing correlation values for larger *L* values.

### D. Energy gap

We now study the energy gaps of structures, i.e., the size of the energy difference between the first suboptimal structure and the energy of the MFE structure. Based on experimental work, it has been reported earlier that the energy gap of some functional RNA is typically larger for natural RNA than for random RNA [58]. Also, that the energy values of the MFE structures are lower in natural RNA as compared to random sequences [59, 60]. Both indicate that natural RNA may be more stable than random RNA.

In Figure 4 we show distributions of the energy gaps (made using kernel density estimation methods). We see the natural and random distribution are overall very similar. Using the Kolmogorov-Smirnov two-sample test to compares the random and natural distributions, we find that for *L* = 20, the *p*-value is 0.55 which is not significant (*>*0.05), suggesting that the natural distribution is statistically identical to the random data. For the other larger lengths, the *p*-values are significant (≪ 0.05), which means that the two comparison distributions are not identical. Having said that, it is clear that the differences in the distributions are small. The KolmogorovSmirnov two-sample test statistic values were all ≤ 0.05, except *L* = 60 for which the test statistic was 0.08. These are all small values on a scale of [0.0, 1.0], indicating the similarity of the distributions.

**FIG. 4.**
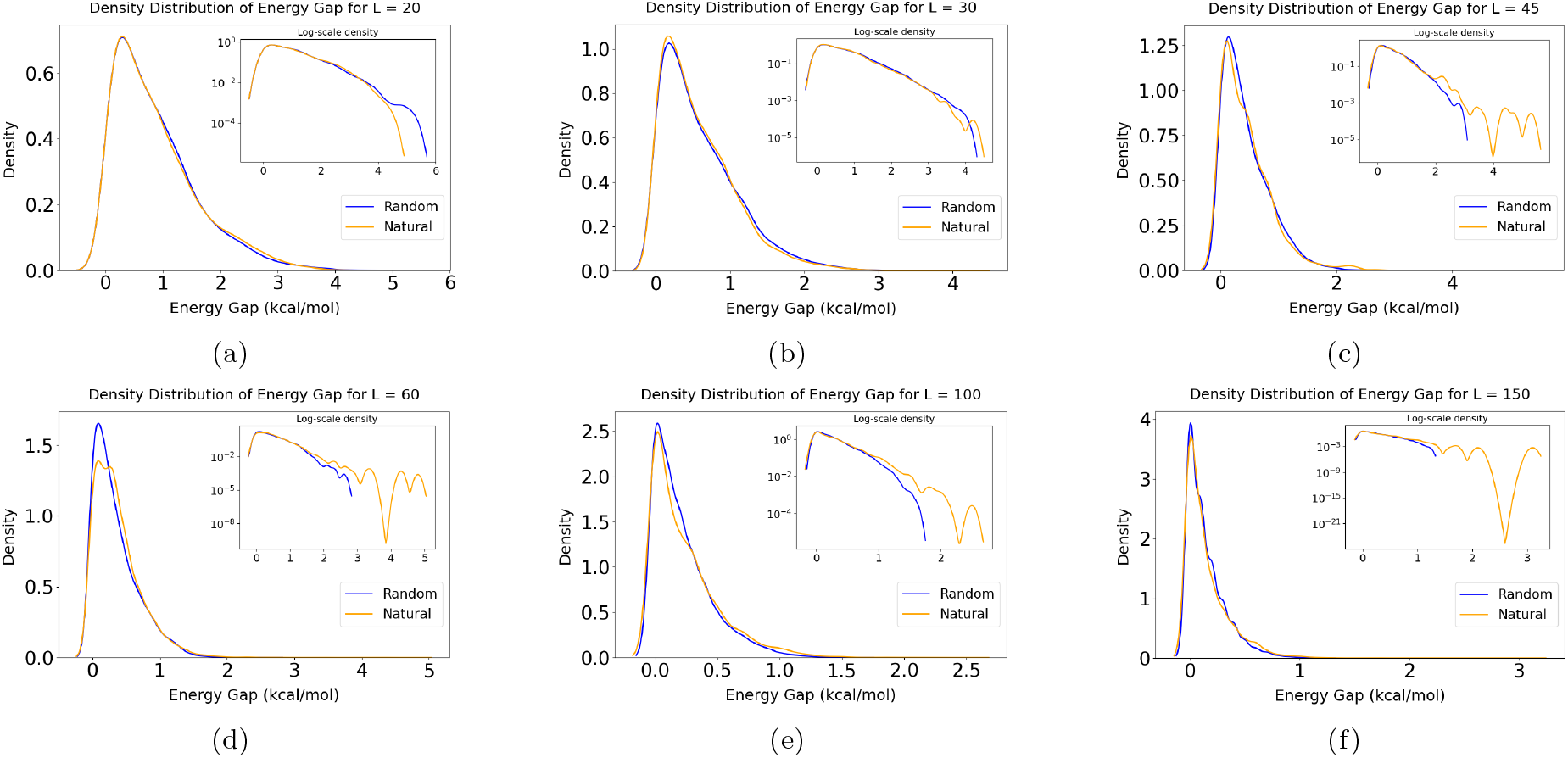
Distributions of energy gaps (kcal/mol) for random and natural RNA. (a) *L* = 20, (b) *L* = 30, (c) *L* = 45, (d) *L* = 60, (e) *L* = 100, and (f) *L* = 150. The natural and random distributions are very similar. Insets show logarithmic *y*-axis, which highlights the differences in the tail of the distributions.

To study the data from another angle and visually appreciate any differences in the right tail of the distributions, the insets in Figure 4 show the same distributions but on a logarithmic *y*-axis. From this perspective, it appears that for *L* = 20, the natural data have *smaller* energy gaps in the tail of the distribution. On the other hand, for *L* ≥ 45, the natural data has an overabundance of large energy-gap structures. The larger energy gaps accord with earlier studies [59, 60], but the smaller gaps for *L* = 20 do not.

### E. Energy gap thresholds

As a different perspective on energy gaps, we investigate the fraction of large gaps in the natural data. In particular, we investigate whether the natural data has a larger or smaller fraction of extreme gap sizes, as compared to the random data. To do this, we take the random sequences data, and for each length *L*, we find the energy gap value threshold (kcal/mol) that separates the top 1% largest gaps from the bottom 99%. Figure 5(a) shows these threshold values. As can be seen, as *L* increases, the energy thresholds decreases. Taking *L* = 20, for example, the value which separates the top 1% largest energy gap values from the rest of the values is about 3.2 kcal/mol, while for *L* = 150 the threshold value is only 0.7 kcal/mol.

**FIG. 5.**
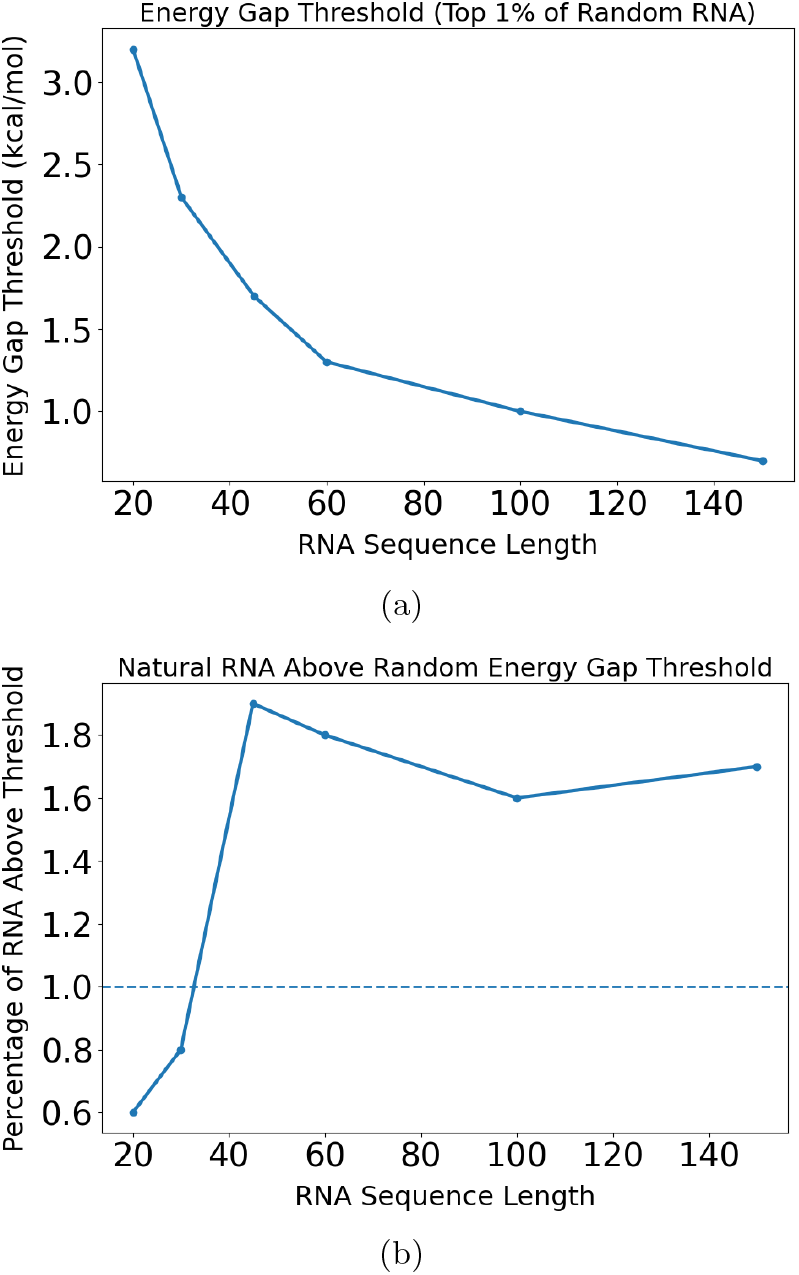
Energy gap thresholds. (a) The energy gap threshold which separates the top 1% (i.e. 99th percentile) for random sequences decreases with increasing length *L*. (b) The fraction of natural sequences with energy gap values above the 1% threshold derived from random RNA. For shorter RNA of *L* = 20, 30 fewer than 1% of natural RNA have energy gap values less than the threshold, but for larger natural RNA of *L* ≥ 45 more than 1% are above the gap threshold.

Next, for each *L*, we find what fraction of natural RNA data have energy gaps larger than the threshold values plotted in panel (a). If the natural and random data are statistically identical, for each *L*, we would expect to see ≈ 1% of the natural data having larger energy gaps than the threshold values in panel (a). Turning to Figure 5(b), we see the fraction of natural RNA data above the random threshold is somewhat close to ≈ 1% for each *L*, but at the same time there is a trend: the fraction increases from below 1% to above 1% as *L* increases. For *L* = 20, only 0.6% of the natural data have gaps larger the 1% threshold from random data, while for *L* = 150, 1.7% of the natural data do.

Interestingly, if we equate larger energy gaps with more stable secondary structures, then these data suggest that longer RNA tend to have a larger fraction of more stable structures, while for small RNA of *L* = 20 or 30, the fraction of natural RNA with large gaps is smaller. This potentially suggests that the very small RNA may be selected for less stability, while the larger RNA are selected for more stability.

### F. Entropy of the Boltzmann distribution

Next, we compare natural and random RNA via the Shannon entropy values of the Boltzmann distribution. For a given sequence, the entropy *H* over the Boltzmann ensemble is given by

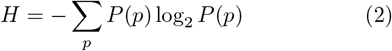

where the summation is over all structures *p* in the Boltzmann ensemble, and *P* (*p*) is the probability of structure *p*. It is well known that 2^*H*^ gives an estimate of the effective number of outcomes of a discrete random variable [61], where *H* is the distribution Shannon entropy in bits.

It follows that if the Boltzmann distribution for some sequence has high entropy, the sequence alternates between many different suboptimal structures. Conversely, if the distribution has low entropy, the sequence alternates between only few structures.

In Figure 6 we see that the distributions of Boltzmann entropy values *H* are similar for RNA with *L* = 20, 30, 45 (mean values given in the figure titles), but differ for the longer RNA *L* = 60, 100, 150, for which the natural entropy values are lower. We also checked the entropy distributions without including the MFE, so as to remove the influence of the energy gap specifically, but the overall picture is the same (not shown). This observation accords with earlier work of Moss [62] who found that natural RNA have lower ensemble diversity than random RNA. In Figure 6, all *p*-values for differences in natural vs. random distributions are significant (*<* 0.05).

**FIG. 6.**
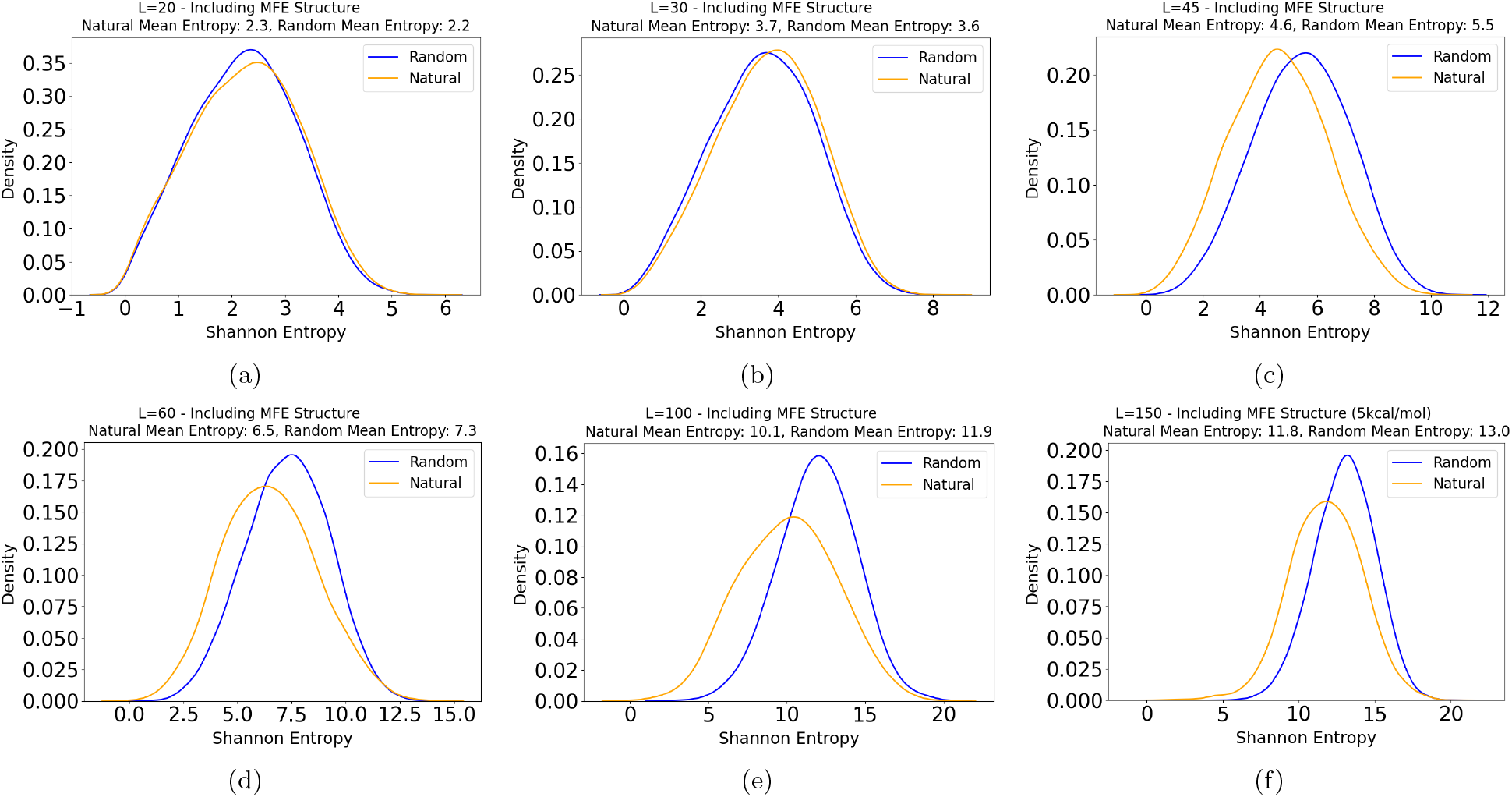
Distributions of Shannon entropy values of the Boltzmann distributions (including the MFE structure). (a) *L* = 20, (b) *L* = 30, (c) *L* = 45, (d) *L* = 60, (e) *L* = 100 (samples of 1000 sequences only), and (f) *L* = 150 (using a cutoff of 5.0 kcal/mol). The distributions are similar for smaller RNA, but differ for larger RNA.

As a brief check that the observed differences in entropy values of larger RNA is not merely a result of sequence composition biases, in Appendix B we also plot the distribution of entropy values for randomly permuted natural sequences. By randomly permuting natural sequences, we can ensure that the composition of nucleotides is the same as the composition of the natural RNA. Such permuting is sometimes used in comparing random and natural RNA [29]. Figure B.8 in the Appendix shows that the permuted and purely random sequence distributions are very similar, and hence the difference between random and natural RNA is not merely due to sequence composition. Also, Figure C.9 shows that for *L* = 60 data, the random and shuffled data are similar.

### G. Ensemble Hamming self-distance

We now make a comparison between the ensembles for random and natural RNA in terms of how much structural variation there is, computed through the Hamming distance.

For a given sequence, we sample two structures from its Boltzmann ensemble, and then find the Hamming distance between the two sampled dot-bracket structures. We call this the *self-distance*, because it is a distance originating from one genotype sequence. We repeat this for many random sequences of length *L*, and compute the mean and standard deviation. We repeat the same procedure for natural data of the same length, and compare the means. This method of Hamming distance quantification follows the work of Novev and Ahnert [63]. If the ensemble is very diverse in terms of having many different structures with relatively high probability, then the mean self-distance will be high. If the ensemble is not diverse, for example if the energy gap is large or there are many structures with high probability but they are very similar to each other, then the self-distance will be low.

Figure 7 shows the mean Hamming self-distance for the different lengths, *L*. Consistent with the analysis of the Shannon entropy, for larger *L* ≥ 45, the mean distance is slightly lower for natural as compared to random. However, for the smaller RNA with *L* = 20 and 30 the mean Hamming distance is very close or even slightly larger for the natural than the random. This suggests that the smaller natural RNA may have more diverse structural variation than the random RNA, contrasting the observations for the larger RNA.

**FIG. 7.**
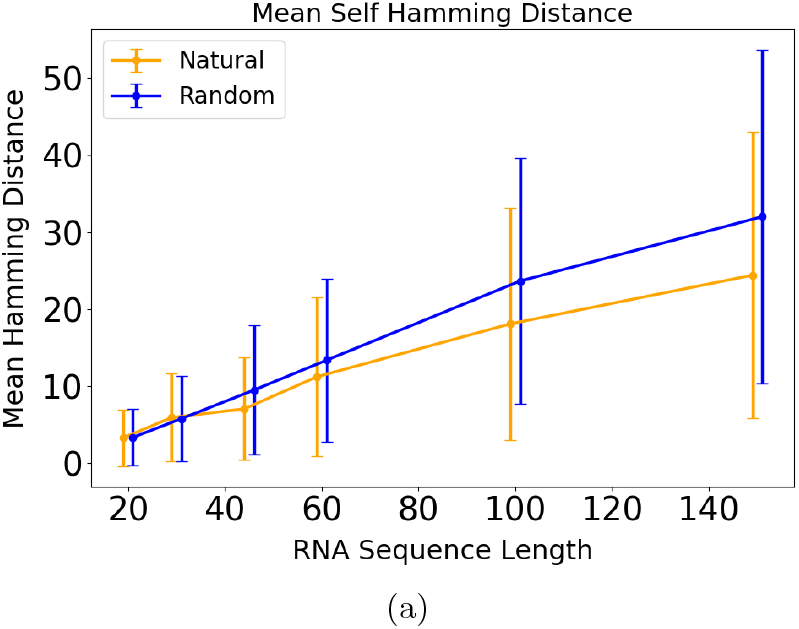
Mean Hamming self-distance of dot-bracket structures. Plots are shown with *x*-axis offset by *±* 1 for visualization purposes. The percentage of the mean (natural) as a fraction of the mean (random) for each length *L* is: 99%, 103%, 74%, 84%, 77%, 76%. For shorter sequences with *L* = 20 and 30, the natural data and random data are almost equal in terms of self-distance. For *L* ≥ 45, the self-distance of the natural data is somewhat smaller than for random data. For each length *L*, data from 1000 random and 1000 natural sequences were used.

### H. Separating RNA types

In our main analysis, the natural dataset combines several different types of RNA. In principle, this could hide type-specific differences between natural and random sequences. As a brief investigation of RNA types, in Appendix D we compare natural data for some common types of ncRNA, namely tRNA, miRNA, rRNA, and snRNA to random RNA. Overall, the distributions and statistics obtained from each RNA type is broadly similar to those from random sequences, thus suggesting that our main claim of a similarity between natural and random RNA ensembles holds also roughly also within RNA types. Having said that, a noteworthy observation is that in our analysis miRNA of *L* = 60 is substantially smaller self-distances than random RNA, implying less structural variety.

This conclusion of overall similarity even when treating RNA groups separately accords with von der Dunk et al. who found that even within RNA types, random and natural RNA secondary structures are similar.

## III. DISCUSSION

Within theoretical biology and evolutionary theory, an interesting question concerns morphospace occupation, that is, the regions of possible form and trait space which are occupied by natural organisms, and those regions which are not. There are various reasons why certain areas of a morphospace may or may not be occupied, including natural selection, developmental biases, historical contingency, complexity [64, 65], and maybe others. In the context of RNA structure morphospaces, previous studies [19, 20, 28, 30] have examined the properties and shapes of RNA secondary structures, but mainly in terms of minimum free energy structure (MFE). To advance the field, here we extend the study by examining energetically suboptimal structures.

Our main conclusion is that for lengths *L* = 20, 30, 45, 60, 100, 150, random and natural non-coding RNA have generally similar Boltzmann ensembles and distribution properties. In other words, the regions of the ensemble morphospace that random and natural noncoding RNA occupy are generally very similar. This suggests that genotype-phenotype map structure itself is a main factor determining the distribution of natural RNA suboptimal secondary structure properties. The small differences we found between natural and random RNA are that natural RNA tend to be more energetically stable and the Boltzmann distribution over the ensemble of suboptimal structures is slightly more concentrated, echoing reports from earlier studies. However, we also found some evidence that shorter RNA of lengths *L* = 20 to 30 are equally or even less stable, and more diverse, than random sequences, in contrast to the longer RNA. This hints, potentially, at some selection for diversity in very small RNAs, contrasting with the observations for longer RNA. This result has been previously studied as RNA plasticity emerging under selection of fluctuating environments [66].

It is known that computational prediction of RNA secondary structures is not 100% accurate [32], and hence this imposes a limitation on our results which employ such algorithms. Having said, it is unlikely that prediction accuracy is a major limitation, because these algorithms have been tested and are reasonably accurate [23], especially for shorter non-coding sequences that we study here [53, 67]. While the cited works examined specifically MFE structure prediction accuracy, not ensemble profile prediction accuracy, we can expect these to be closely related. We are not aware of any previous work which explicitly tests the accuracy of the Vienna RNA package predictions of suboptimal structures, but this would be valuable for future work. Additionally, the similarity between natural and random RNA structures in terms of other traits has been independently established via comparing to experimental and consensus structures [19, 20, 30].

Another limitation of the current study is that we examined only short sequences of *L* ≤ 150, while natural RNA lengths extend into the 1000s of bases. However, earlier works [29, 68] have studied larger RNA and found similarities between natural and random RNA, so it is unlikely that our results are only relevant to shorter RNA. We leave studying longer RNA for future work.

An main limitation of our study is that the Boltzmann distribution is relevant when a molecule is assumed to be in equilibrium, but the cell of an organism is not in equilibrium, but is instead a complex environment of different chemical and dynamic processes. It follows that the Boltzmann probabilities of suboptimal structures which we compute here may not be entirely relevant to the molecules in the cell. Due to this limitation, our work (and related earlier studies) act as a first order approximation to understanding the distribution over suboptimal structures. Future work could include a comparison of natural and random RNA sequences with kinetic folding simulations (e.g., Gillespie-based or coarse-grained molecular dynamics) to give a more detailed picture.

It would be very interesting to study RNA tertiary structures, but at present computational tools for predictions of full RNA forms are lacking [33]. This can be left for future work. Additionally, while several earlier papers and this work have studied properties of natural RNA for given fixed lengths, it would be interesting to study the distribution of natural RNA lengths, especially in terms of whether there are biophysical factors that influence or predict the distribution of natural lengths.

Another route for further work would be to study the combined shapes for longer chains of shapes. Here we used the first 5 shapes, but in the tail of the ensemble there may be more variety and subtle differences. However, longer chains pose a challenge because they increase sample space sizes exponentially, and hence increase the amount of random sampling and natural data required to obtain reliable frequency statistics

As discussed in the Introduction, it is by no means obvious or trivial that natural functional non-coding RNA should have secondary structures (and suboptimal ensemble profiles) similar to typical random RNA. This is because in biochemistry it is well known that structure and function are closely linked, and hence the structures of these RNA are important for their functions, and also under selection. So, how can we rationalize this similarity? The first possible explanation is the *arrival of the frequent* [69] hypothesis. This hypothesis says that in genotype-phenotype maps with strong bias, the order of the appearance of phenotypes, upon random mutation, has a strong effect on which phenotypes fix. In fact, the bias can dominate over the influence of fitness; see also [70, 71] and earlier work on bias is the introduction of variation [72, 73]. The close agreement between random and natural ncRNA could be explained by saying that once a secondary structure is found that is ‘good enough’, selection mainly works by further refining parts of the sequence for function, rather than significantly altering the structure [20]. Indeed, there may be many such structures which are good enough [30, 74] and so from these the most likely are chosen. Another possible explanation, discussed in ref. [29], and which is more speculative, is that there may be some connections between structures with high probability and fitness. For example, robustness to genetic mutations is a property which has a fitness benefit, and yet robustness is positively correlated with phenotype probability [75, 76]. Hence, it is not beyond the realm of possibility that high-probability structures appear in the natural data not only because of their higher chance of appearance, but due to some selection effects. More speculatively, it has been argued from algorithmic information theory and several concrete examples that there may be a general connection between optima and high-probability structures [77, 78]. Again, this would point to a possible connection between fitness and probability.

Looking to future work, it would be valuable to investigate other biomolecules in terms of morphospace occupation, as well as shapes and patterns higher up in the levels of biological organization [65]. Further, more work is needed in understanding and explaining the close similarity between random and natural RNA in terms of morphospace occupation.

## Data and code

The code used for this study is available from

github.com/Hibaa-K/comparing-rna-boltzmann-ensembles.

## Acknowledgments

This project has been partially supported by the Kuwait Foundation for the Advancement of Sciences (KFAS), grant number PN24-12SL-2290.

## Appendix A Methods

### 1. RNA sequences

For the natural RNA, we took all available sequences of lengths *L* = 20, 30, 45, 60, 100, 150 from the RNAcentral database [25]. The number of natural sequences for each length is 12,438 (*L* = 20), 47,975 (*L* = 30), 13,541 (*L* = 45, see below), 20,897 (*L* = 60), 25,533 (*L* = 100) and 15,494 (*L* = 150).

We cleaned the data by removing any duplicates and filtering out sequences containing non-standard nucleotides, keeping only those composed of A, U (T), C, and G. For the random sequence data, a pseudo-random number generator was employed to make random sequences of the same lengths.

For *L* = 45, there was an over abundance of just one particular RNA, namely hammerhead ribozyme sequences from Schistosoma species (58,400 from a data set of around 76,000 sequences). Therefore, to reduce this sampling bias, all sequences of Schistosoma species, specifically the hammerhead ribozyme type, were excluded. This was done in addition to the cleaning and filtering that was performed for the other datasets of RNA sequences.

### 2. Structures and shapes

The MFE secondary structures and suboptimal structures were predicted computationally via energy minimization using the well-known RNA Vienna package [23], with default parameters (we use *T* = 37 degrees centigrade). The algorithm gives predicted energy values for the MFE and suboptimal structures to within a given threshold given in kcal/mol.

Suboptimal shapes were calculated using the RNAsubopt function of the Vienna package, see www.tbi.univie.ac.at/RNA/RNAsubopt.1.html

RNA secondary structures can be abstracted in standard dot-bracket notation, where brackets denote bonds, and dots denote unbonded pairs. To obtain coarse-grained abstract shapes [57] of differing levels we used the RNAshapes tool available at https://bibiserv.cebitec.uni-bielefeld.de/rna-shapes and the Bioconda rna-shapes package available at https://anaconda.org/bioconda/rnashapes. The option to allow single-bonded pairs was selected, to accommodate the Vienna folded structures which can contain these.

### 3. Outlier analysis

To define outliers for energy gap distributions, we used the standard criterion that any sample with size larger than *Q*3 + 1.5*IQR* is declared an outlier, where Q3 is the third quartile, and the IQR is the inter-quartile range. We did not use the more common criterion of 3 standard deviations from the mean, because the distributions of energy gaps are not normal. The values of Q3 and IQR were derived from the natural RNA data.

## Appendix B Shannon Entropy with Shuffled Natural RNA

As a brief check that the differences in entropy distributions observed between natural and random is not simply due to the natural sequences having different sequence compositions (i.e., in terms of CG content), we also studied the distributions of entropy value for randomly shuffled/permuted sequences. These sequences would have the same sequence compositions as the natural RNA sequences. As seen in Figure B.8, the shuffled and purely random sequences are similar. This shows that the differences between the random and natural RNA entropy values are not merely due to sequence composition factors.

## Appendix C Choice of energy threshold cutoff

### 1. Tradeoff

As discussed in the main text, we chose to use an energy threshold of 10.0 kcal/mol in the analysis of the Boltzmann distribution for *L* ≤ 100, and 5.0 kcal/mol for *L* = 150. The choice of threshold is a tradeoff between accuracy and computational efficiency: larger thresholds are (potentially) more accurate due to including more suboptimal structures, while smaller thresholds are more computationally efficient. We say “potentially” more accurate, because eventually, increasing the cutoff value will include only structures with negligible contributions to the ensemble.

The choice of a smaller threshold of 5.0 kcal/mol for *L* = 150 is guided by computational tractability. Larger structures have many more suboptimal structures, for a given threshold. Using 10.0 kcal/mol proved overly computationally demanding for our dataset. At the same time, this means that the results for *L* = 150 are likely to be less accurate, compared to the analysis from the other lengths.

Note also that the choice of cutoff does not impinge on all of our results, for example the distribution in the size of energy gaps is not sensitive to the choice of threshold.

**FIG. B.8.**
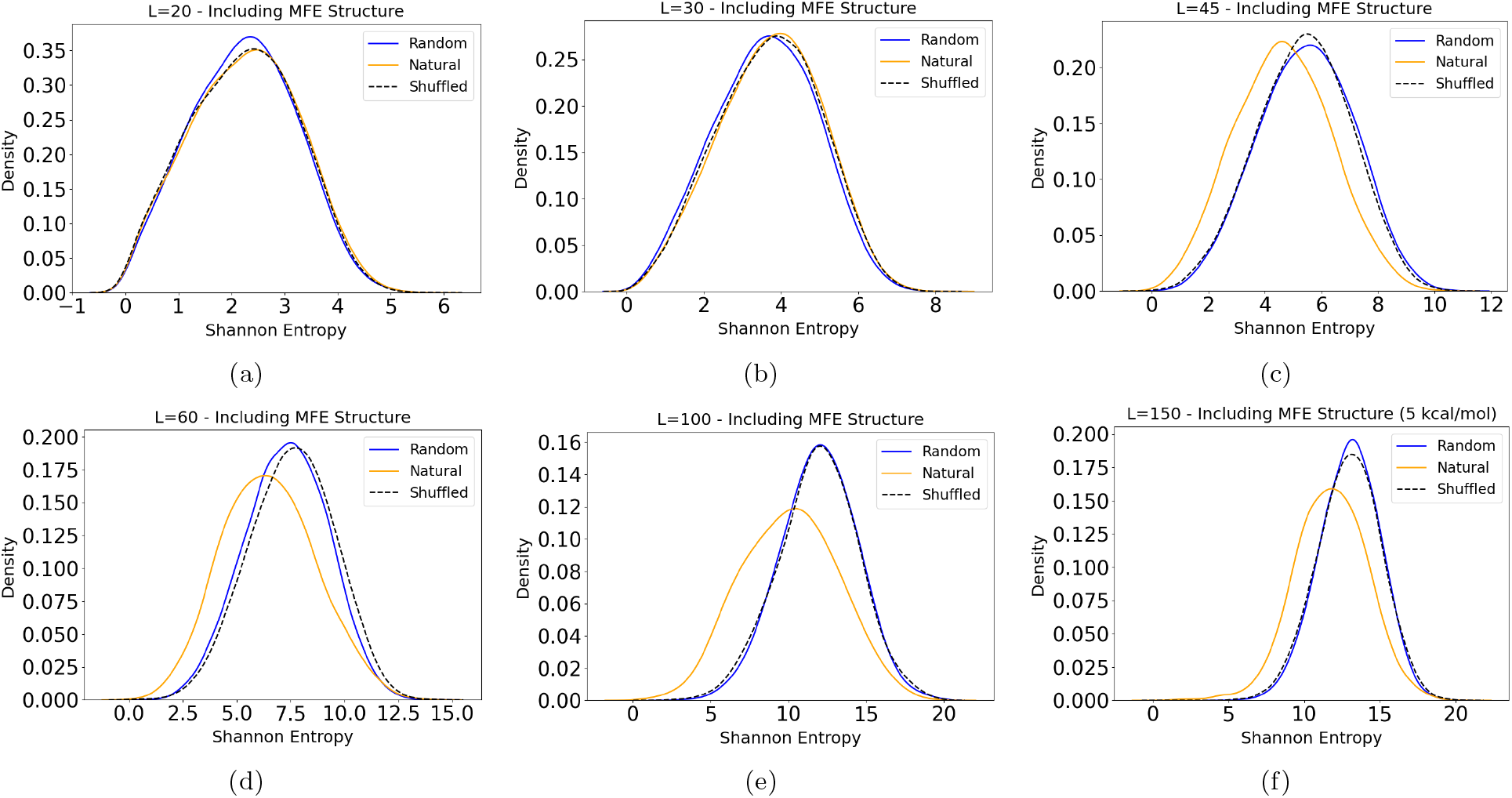
Distributions of Shannon entropy values for random, natural, and shuffled/permuted. Data includes the MFE structure for natural, random, and shuffled/permuted natural RNA data. Except for the very short RNA, the randomly permuted and purely random sequences show similar distributions.

### 2. Empirical check

As a brief check that the choice of 10.0 kcal/mol is a reasonable tradeoff value, we look at the mean Hamming self-distance for natural data of *L* = 60 for different choices threshold (using a sample of 5000 natural sequences). Hamming distance is likely to be sensitive to energy threshold, because higher thresholds presumably allow for greater diversity in structures, and hence larger mean Hamming distances. This sensitivity implies that it is a good choice for experimenting with different cutoff values.

Performing the computations for the *L* = 60 data yields the following values (mean and standard deviation):

~~~
Calculating Mean Hamming Self Distance For L=60 Natural RNA (5000 seqs sample):
- 3.0 kcal/mol: 9.7 (+/- 9.9)
- 5.0 kcal/mol: 10.8 (+/- 10.1)
- 10.0 kcal/mol: 11.2 (+/- 10.1)
- 12.0 kcal/mol: 11.1 (+/- 10.0)
~~~

As can be seen, when moving from 3.0 kcal/mol to 5.0 kcal/mol, there is a substantial change in the mean Hamming distance (9.7 vs. 10.8), suggesting that using 3.0 would be too small, and would negatively impact accuracy. However, the difference in mean values when using 5.0 kcal/mol and 10.0 kcal/mol is modest (10.8 vs. 11.2), and the difference between mean values when using 10.0 kcal/mol and 12.0 kcal/mol is very small (11.2 vs. 11.1). This suggests that 10.0 kcal/mol is a decent cut off value, and that using larger and larger values would have a relatively small impact on the quantitative results of our study.

**FIG. C.9.**
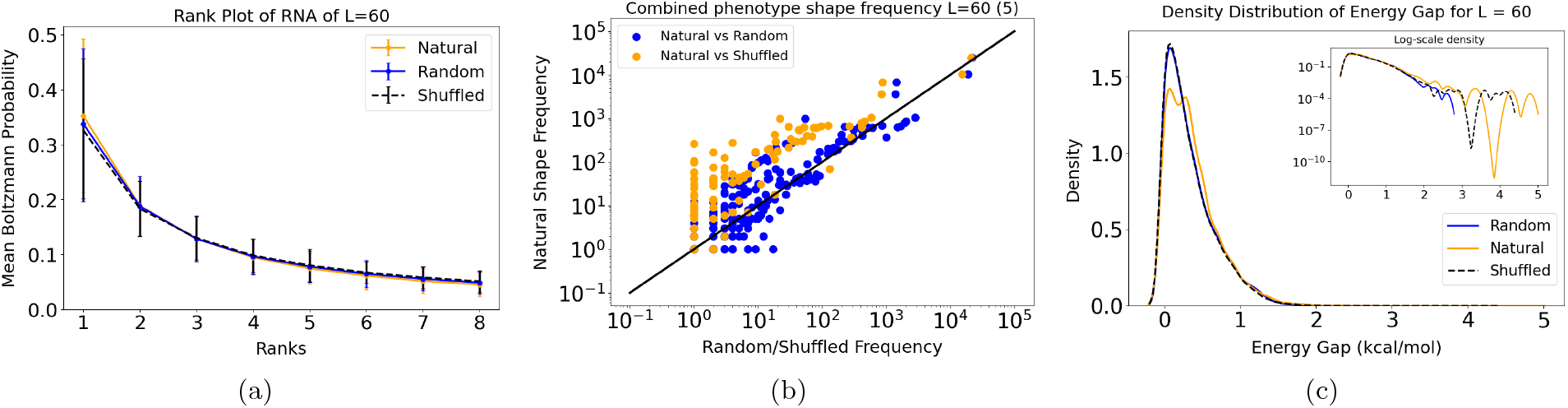
Comparing shuffled/permuted natural RNA data for *L* = 60 to natural and random sequences. (a) The rank plot of mean Boltzmann probabilities is similar for all three of shuffled, natural, and random data. (b) Correlation plots of combined shape frequencies show high correlation for both natural vs. random, and natural vs. shuffled. (c) The distribution of energy gaps for random and shuffled sequences is very similar for the main body of the distribution, but the tail fluctuates more for the shuffled than the random sequences (somewhat similarly to the natural data). Overall for *L* = 60, the shuffled sequences are similar to the random.

## Appendix D Analyzing RNA types

In the main text, the natural data combine several different types of RNA. In principle, this could hide type-specific differences between natural and and random sequences. Here we study natural data separated by RNA type.

### 1. RNA types: Large energy gaps

Looking more closely at the specific RNA with larger energy gaps, for each *L*, we find the sequences which are considered outliers in terms of large gaps (Methods, Appendix A). In Appendix E we give the details of which natural RNA had energy gaps deemed to be outliers and summarize the main findings here: For each length *L*, we give the three most common types of RNA as labeled in the database

- *L* = 20 miRNA (25%); unknown (22%); snRNA (20%)
- *L* = 30 piRNA (71%); small RNA (12%); rRNA (6%)
- *L* = 45 ribozyme (63%); unknown (18%); tRNA (6%)
- *L* = 60 tRNA (49%); miRNA (28%); rRNA (10%)
- *L* = 100 SRP RNA (23%); tRNA (20%); miRNA (19%)
- *L* = 150 SRP RNA (41%); rRNA (11%); miRNA (8%)

We see that miRNA, tRNA, rRNA appear frequently in these collections of RNA with larger energy gaps. Given that these are essential RNAs for regulation of gene expression through binding, the enhanced stability might be due to their regulatory functions being highly dependent on their structural integrity.

### 2. RNA types: Energy gaps

As a brief investigation of RNA types, we took natural data for *L* = 60, for four common types of RNA: tRNA, miRNA, rRNA, and snRNA. Figure D.10 shows the distributions of energy gaps for these four types, as compared to random sequences. In all cases, the *p*-value is ≪0.05 indicating that none of the distributions are statistically identical to random samples. More importantly for our purposes however, is that the distributions are each generally very similar to the random distributions. More quantitatively, the Kolmogorov-Smirnov test statistics are all small, with 0.11 for tRNA, 0.11 for miRNA, 0.07 for rRNA, and 0.08 for snRNA.

**FIG. D.10.**
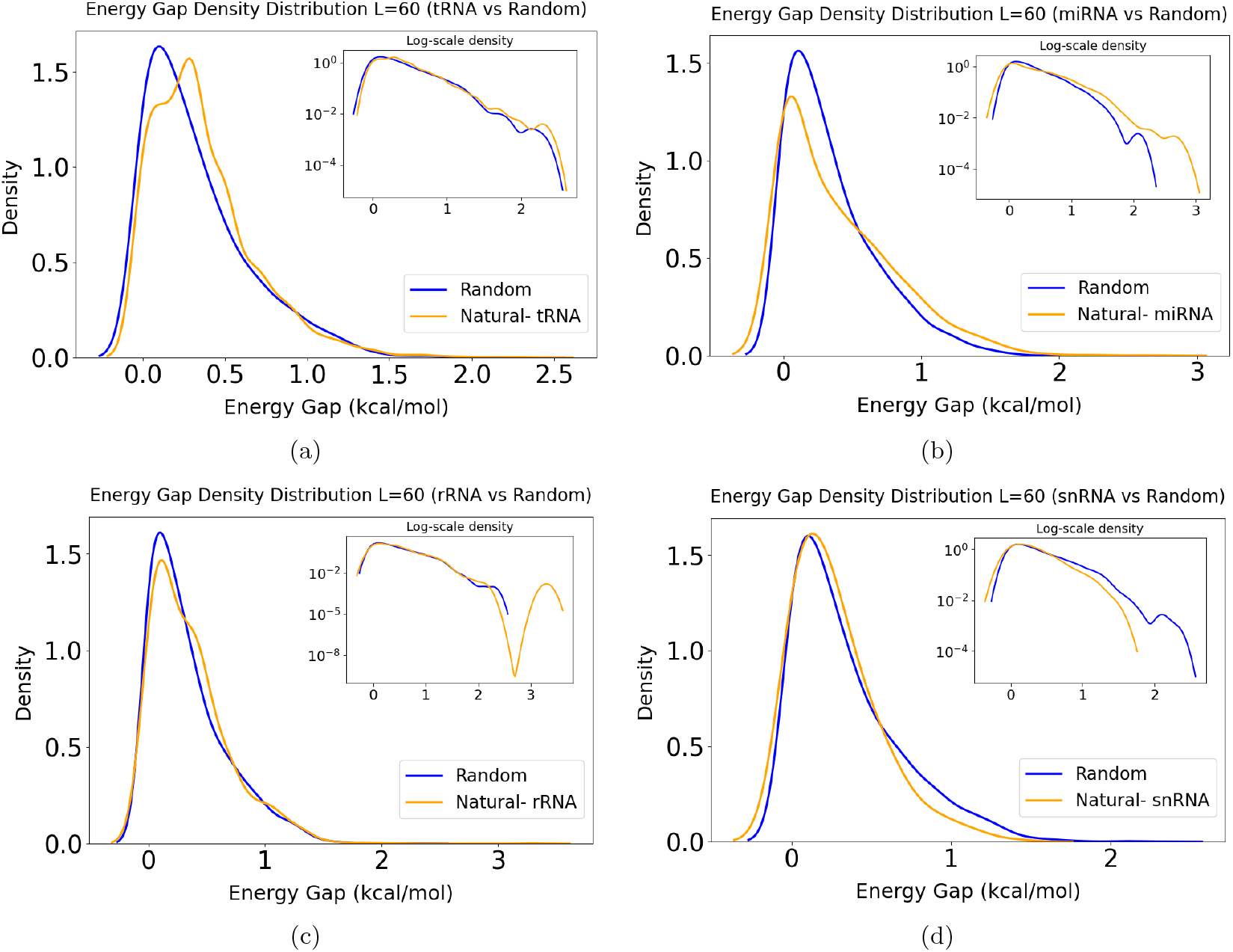
Distribution of energy gaps for four common natural RNA types vs. random with *L* = 60. In each case, a sample of 5,000 random sequences were used. (a) tRNA (12,875 seqs.), (b) miRNA (3,299 seqs.), (c) rRNA (2,334 seqs.), and (d) snRNA (390 seqs.).

Visual inspection of the distributions shows that tRNA and the random data are very similar indeed, while miRNA shows a noticeable different in the form of the distribution, and in the tail. It is interesting that snRNA shows a similar distribution in the body, but the tail appears different, with snRNA appearing to have smaller energy gaps that the random sequences in the tail.

### 3. RNA types: Hamming self-distance

Continuing from the previous subsection, we make a brief study of the mean Hamming self-distance for natural RNA, where we have separated the RNA into classes, choosing four common types of RNA: tRNA, miRNA, rRNA, snRNA. We used a sample of 1000 sequences for each type except for snRNA which has 390 sequences. All sequences have *L* = 60:

~~~
Mean self hamming distance using 10 kcal/mol:
tRNA: 12.5 (+/- 10.62)
miRNA: 5.4 (+/- 4.63)
rRNA: 11.9 (+/- 9.53)
snRNA: 12.9 (+/- 9.57)
~~~

For comparison, the random sequences of *L* = 60 had a mean Hamming self-distance of 13.4. Therefore, trna and snRNA have slightly lower mean values compared to random sequences, but not substantially so. rRNA has a somewhat smaller mean than random sequences, but miRNA is a clear outlier with a far smaller mean than for random sequences.

## Appendix E Natural data - energy gap outliers

The detailed records of the outlier fractions and values in relation to (see Figure 4) in the main text are given below:

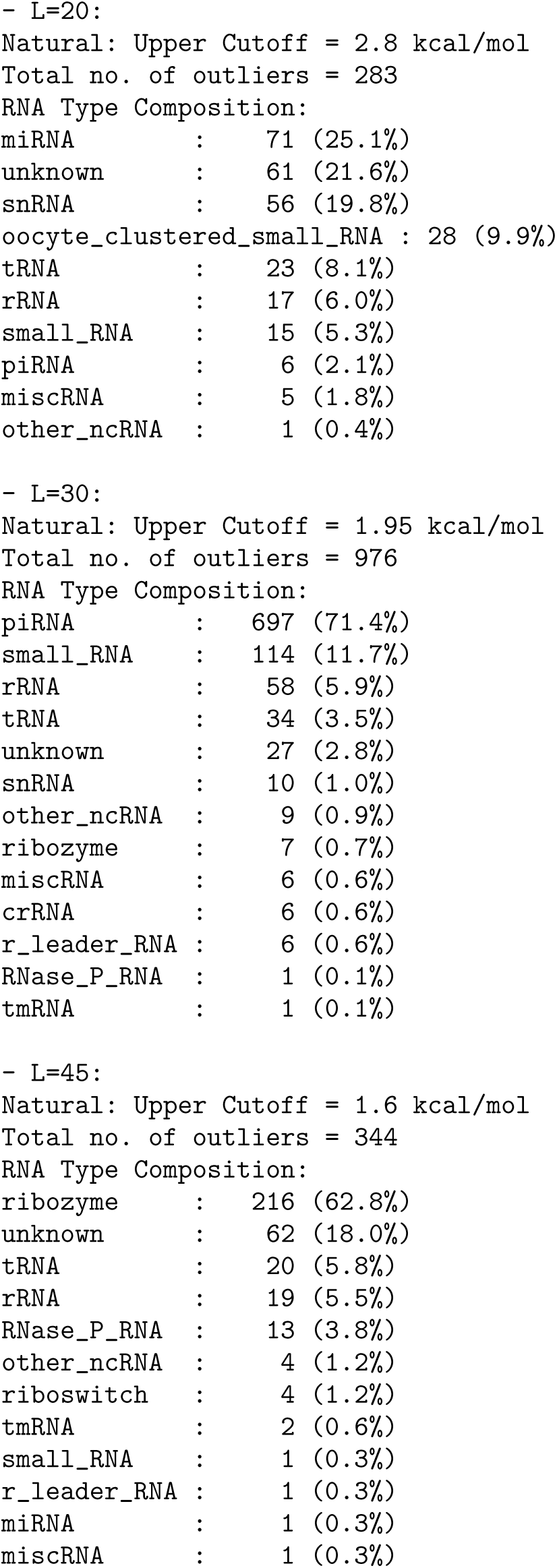
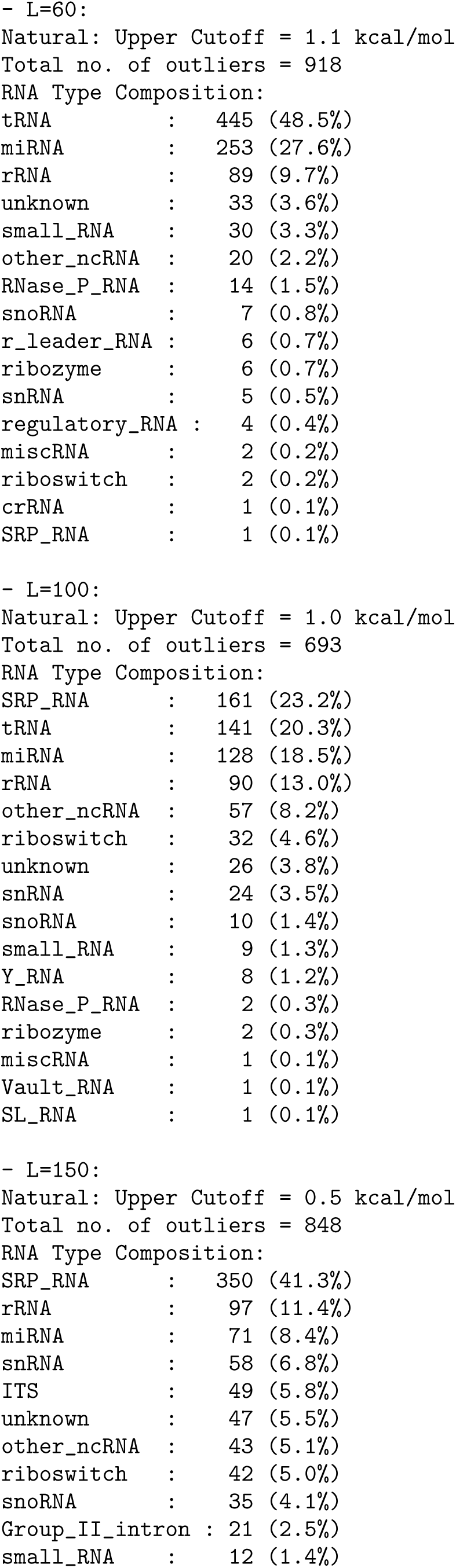
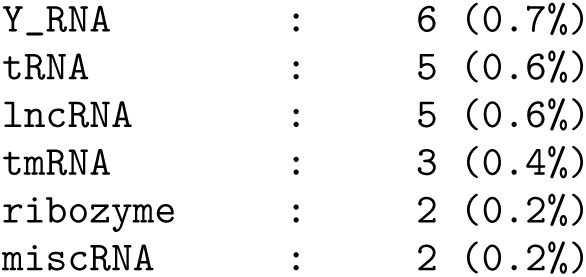

